# Improved *de novo* Genome Assembly: Linked-Read Sequencing Combined with Optical Mapping Produce a High Quality Mammalian Genome at Relatively Low Cost

**DOI:** 10.1101/128348

**Authors:** DW Mohr, A Naguib, NI Weisenfeld, V Kumar, P Shah, DM Church, D Jaffe, AF Scott

## Abstract

Current short-read methods have come to dominate genome sequencing because they are cost-effective, rapid, and accurate. However, short reads are most applicable when data can be aligned to a known reference. Two new methods for *de novo* assembly are linked-reads and restriction-site labeled optical maps. We combined commercial applications of these technologies for genome assembly of an endangered mammal, the Hawaiian Monk seal.

We show that the linked-reads produced with 10X Genomics Chromium chemistry and assembled with Supernova v1.1 software produced scaffolds with an N50 of 22.23 Mbp with the longest individual scaffold of 84.06 Mbp. When combined with Bionano Genomics optical maps using Bionano RefAligner, the scaffold N50 increased to 29.65 Mbp for a total of 170 hybrid scaffolds, the longest of which was 84.78 Mbp. These results were 161X and 215X, respectively, improved over DISCOVAR *de novo* assemblies. The quality of the scaffolds was assessed using conserved synteny analysis of both the DNA sequence and predicted seal proteins relative to the genomes of humans and other species. We found large blocks of conserved synteny suggesting that the hybrid scaffolds were high quality. An inversion in one scaffold complementary to human chromosome 6 was found and confirmed by optical maps.

The complementarity of linked-reads and optical maps is likely to make the production of high quality genomes more routine and economical and, by doing so, significantly improve our understanding of comparative genome biology.

## INTRODUCTION

High quality non-human genomes, especially mammalian genomes, are needed for a variety of reasons including 1) better constraining the limits of allowable nucleotide variation (a useful tool for identifying potentially causative mutations in human disease), 2) the identification of conserved protein and regulatory regions that may explain the morphological or physiological characteristics of different species, 3) establishing the correct relationship of SNPs to genes for association studies, 4) improving our understanding of evolutionary relatedness and mechanisms, and 5) aiding efforts for species conservation and management. The need for fast, economical and high quality genome data for *de novo* genome assembly of rare and endangered species will become especially important as habitat loss, climate change and human impacts accelerate in the 21^st^ century.

Although next generation sequencing has allowed fast and accurate sequencing, to date, the largest hurdle to genome assembly has been the difficulty and cost in obtaining long contiguous sequence scaffolds from short reads. The human genome and those of many model organisms were assembled using methods where large DNA molecules were cloned (e.g., BACs, cosmids, etc.), employed long DNA sequencing technologies (e.g., PacBio), or used augmented short-reads approaches such as mate-pair libraries. Recently, new single-molecule chemistries and analytic methods have been developed to extend these approaches. Here, we have evaluated two technologies for assembling a *de novo* mammalian genome of the Hawaiian monk seal (*Neomonarchus schauinslandi*), an endangered species endemic to the Hawaiian islands, using techniques that interrogate individual long DNA molecules and have compared these to the short-read based DISCOVAR method (Weisenfeld et al., 2014) which uses overlapping paired-end (PE) Illumina reads. The first method, the 10X Genomics (10XG) Chromium chemistry, incorporates unique molecular indexes (UMI) into single long DNAs and assembles these into gapped scaffolds, based on shared UMIs, using the Supernova software tool (Weisenfeld et al., 2016). The second method used optical maps from Bionano Genomics (Bionano) both as a quality assessment for the Chromium sequences and as a tool to merge the Supernova scaffolds into longer hybrid assemblies.

## METHODS

### Sample

Blood (∼ 10 ml) was collected in EDTA vacutainers during surgery of an adult male seal, shipped on ice packs and processed within two days of collection. The viability of the cells was assessed by trypan blue exclusion. 1 ml of whole blood was stored in LN2 while 9 ml (1.85x106 cells/ml) were used for lymphocyte separation and subsequent DNA isolation for optical mapping.

### DISCOVAR libraries

1 ug of DNA was used to prepare PCR-free Illumina libraries of ∼ 450 bp mean insert size as described on the Broad Institute website (https://software.broadinstitute.org/software/discovar/blog/?page_id=375). We generated 703 million 250bp paired-end reads on an Illuina HiSeq2500 for the PCR free library, for a read depth of approximately 75x based on estimated genome size. Analysis was done using DISCOVAR de novo v52488 using default parameters (Weisenfeld et al., 2014).

### 10X Genomics Chromium

DNA was isolated using MagAttract (Qiagen) and the molecular weight assayed by pulsed field gel electrophoresis. HMW gDNA concentration was quantitated using a Qubit Fluorometer, diluted to 1.25 ng/ul in TE, and denatured following manufacturers recommendations. Denatured gDNA was added to the reaction master mix and combined with the Chromium bead-attached primer library and emulsification oil on a Chromium Genome Chip. Library preparation was completed following the manufacturer’s protocol (Chromium Genome v1, PN-120229). Sequencing-ready libraries were quantified by qPCR (KAPA) and their sizes assayed by Bioanalyzer (Agilent) electrophoresis.

The library was sequenced (151 X 9 X 151) using two HiSeq 2500 Rapid flow cells to generate 975 M reads with a mean read length of 139 bp after trimming. The read 2 Q30 was 87.93% and the weighted mean molecule size was calculated as 92.33 kb. Mean read depth was approximately 61X. The sequence was analyzed using Supernova software (10X Genomics; Weisenfeld et al., 2016) which demultiplexed the Chromium molecular indexes, converted the sequences to fastq files and built a graph-based assembly. The assemblies, which diverge at “megabubbles,” consist of two “pseudohaplotypes.” The sequence data were originally analyzed using Supernova 1.0 and then repeated using v1.1 which estimates gap sizes rather than introducing an arbitrary value of 100 Ns. As noted above, the Supernova scaffolds were used by the Bionano Hybrid Scaffold tool to create sequence assemblies.

### BioNano Genomics

Optical mapping of large DNA (Xiao et al., 2007) incorporates fluorescent nucleotides at sequence specific sites, visualizes the labeled molecules and aligns these to each other and to a DNA scaffold (Shelton et al., 2015). Lymphocytes were processed following the IrysPrep Kit for human blood with minor modifications. Briefly, PBMCs were spun and resuspended in Cell Suspension Buffer and embedded in 0.6% agarose (plug lysis kit, BioRad). The agarose plugs were treated with Puregene Proteinase K (Qiagen) in lysis buffer (Bionano Genomics) overnight at 50° C and shipped for subsequent processing (S. Brown, KSU). High Molecular Weight (HMW) DNA was recovered by treating the plugs with Gelase (Epicenter), followed by drop dialysis to remove simple carbohydrates. HMW DNA was treated with Nt. BspQI nicking endonuclease (New England Biolabs) and fluorescent nucleotides incorporated by nick translation (IrysPrep Labeling-NLRS protocol, Bionano). Labeled DNA was imaged on the Irys platform (Bionano) and more than 234,000 Mb of image data were collected with a minimum molecule length of 150 kb.

### Bionano Genomics Analysis

Haplotype aware *de novo* assembly was done using the Bionano Genomics RefAligner Assembler (version 5122) software based on the overlap-layout-consensus paradigm (Xiao et al., 2015; Xiao et al., 2007; Anantharaman et al., 2001; Valouev et al., 2006). To build the overlap-layout-consensus graph, first the single molecule optical maps underwent a pairwise alignment where each molecule was aligned to every other molecule. Pairwise alignments generated using the Bionano Genomics RefAligner were used as input to the layout and consensus assembly stage where a draft assembly was built. We used a P value threshold of 1e-10 for the initial assembly and a minimum molecule length of 150kb. Next we refined the draft assembly using a P value cut off of 1e-11. Refinement of the assembly corrected errors and trimmed and split contigs if errors were found. Our first set of refined contigs were further improved using five iterations of extension and merging during which all contigs were aligned to each other to check for overlap or redundancy; the P value threshold used was 1e-15. Map extension was done by aligning all input molecules to the refined maps. When a set number of molecules extended past the end of a contig, they were combined into a consensus and added to the end of the contig. This increased the size of the contigs. A final refinement step was performed on the maps after the iterative cycles of extension and merging to produce a more accurate and haplotype separated final consensus map. Haplotype separated maps were built by aligning molecules to the genome maps and clustering them into two alleles. When the reported difference between the two alleles was large enough, two haplotype separate maps were generated.

### Hybrid scaffolding

Hybrid scaffolding was performed on the haplotype separated genome maps and the Supernova pseudohaplotype scaffolds. The first step involved in silico digesting the Supernova sequence assembly using the Nt.BspQI recognition motif to generate map coordinates. Next the sequence scaffolds were aligned with the genome maps to flag alignments that were concordant and were fed into the iterative merging stage. The P value threshold used for the initial alignment was 1e-10 while merge was set at 1e-11. After the iterative merge stage the hybrid scaffolds were generated and aligned to the original sequence scaffolds. Finally, we exported back from genome map coordinates to FASTA format along with an AGP file (https://www.ncbi.nlm.nih.gov/assembly/agp/AGP_Specification/)” www.ncbi.nlm.nih.gov/assembly/agp/AGP_Specification/) that tracked had been merged.

### Scaffold Quality Assessment

QUAST (Gurevich et al., 2013) was used to generate N50 plots and comparative metrics for the assembly methods. BUSCO (Benchmarking Universal Single Copy Orthologs), a tool for assessing genome completeness (Simão et al., 2015), was run on the hybrid pseudo-reference using 3,023 vertebrate-specific single copy orthologs in genome assembly assessment mode.

### Conserved Synteny

The quality of the Bionano/Supernova hybrid scaffolds was determined in various ways. First, we chose one pseudohaplotype of all 170 unique scaffolds generated from the Bionano Hybrid Scaffold tool as a pseudo-reference and performed TBLASTN (e.g., Jun et al., 2009) to translate the seal sequences for alignment to both human and dog protein databases. The position of matching proteins from known coding genes in the seal pseudo-reference was manually reviewed in IGV (Thorvaldsdottir et al., 2013). We compared the regions of conserved syntenic genes primarily to the human genome because it is best annotated and, in some instances, to dog, cat and other mammals for which longer scaffolds were available.

### DNA alignment

Conserved syntenic gene order was confirmed using nucmer (Kurtz et al., 2004) to align the seal hybrid pseudo-reference to human (GRCh37/hg19), with a minimum cluster length of 20 and a minimum match length of 50. Plots were generated for all matches greater than 1kb.

### Analysis software

Supplement 1 lists the specific software and commands used for analysis.

## RESULTS

Both 10XG and Bionano software are haplotype aware. For this study we chose Supernova “pseudohaplotype” 1 for analysis. No major differences in length were noted between haplotypes, as might be expected for a species with a population size of about 1400 animals and with reported low heterozygosity (Schultz et al., 2009)

A summary of results is shown in Table 1. The 250bp PE DISCOVAR sequencing produced over 437,230 reads with a scaffold N50 of 137,851 bp. In contrast, the 10X Chromium sequencing assembled with Supernova v1.0 had an N50 of nearly 14.6 Mbp, an improvement of over 100 fold. When Supernova v1.1 was used the N50 increased to 22.23 Mbp. The Bionano optical maps significantly improved overall scaffold length and decreased the total number of scaffolds from 203 to 170.

**Table 1.** A comparison of Bionano Irys hybrid scaffolds with 10XG Chromium scaffolds assembled with Supernova v.1.0 (col 2) and v.1.1 (col 3). Col 4 is the 10XG Supernova v1.1 data alone and col 5 is data assembled with DISCOVAR. In addition, 7,696 mostly short scaffolds were unscaffolded with the Irys assembler; their length totaled 40.4 Mbp.

| Assembly | BNG_10X_hybrid<br>Supernova 1.0 | BNG_10X_hybrid<br>Supernova 1.1 | 10X Supernova<br>v1.1 | DISCOVAR |
| --- | --- | --- | --- | --- |
| Total scaffolds | 216 | 170 | 7,932 | 437,230 |
| Largest scaffold | 70.55 Mbp | 84.77 Mbp | 84.06 Mbp | 1.10 Mbp |
| N50 | 20.87 Mbp | 29.65 Mbp | 22.23 Mbp | 137,851 |
| N50 Fold<br>Improvement | 105 | **215** | 161 | 1 |
| L50 | 49 | 26 | 36 | 5100 |
| # N's per 100<br>kbp | 254.65 | 2,183.45 | 2,195.29 | 96.06 |
| Est. total<br>assembly length | 2,318 Mbp | 2,360 Mbp | 2,400 Mbp | 2,462 Mbp |

Figure 1 shows a QUAST (Gurevich et al., 2013) plot of scaffold size distributions. The QUAST statistics estimated the total assembly length from 2.32-2.46 Gbp, which is similar to that of other carnivores. The total number of N’s increased (Table 1) between Supernova v1.0 and v1.1, as expected, due to the improved gap estimation algorithm. The change improved alignment to the optical maps, by more accurately spacing *BspQ*I sites with respect to the sequence scaffolds. Figure 2 shows the corresponding improvement in Bionano confidence scores with Supernova v1.1. Figure 3a shows, in IrysView, an example of a long (∼ 450 kbp) region of high concordance between the optical maps and the Supernova v1.1 scaffold while Fig. 3b shows a region where Supernova may have added larger gaps than appear in the Bionano maps. Figure 4 is an example of where a Bionano map merged two Supernova sequence scaffolds. This example was further studied by conserved synteny analysis.

**Figure 1.**
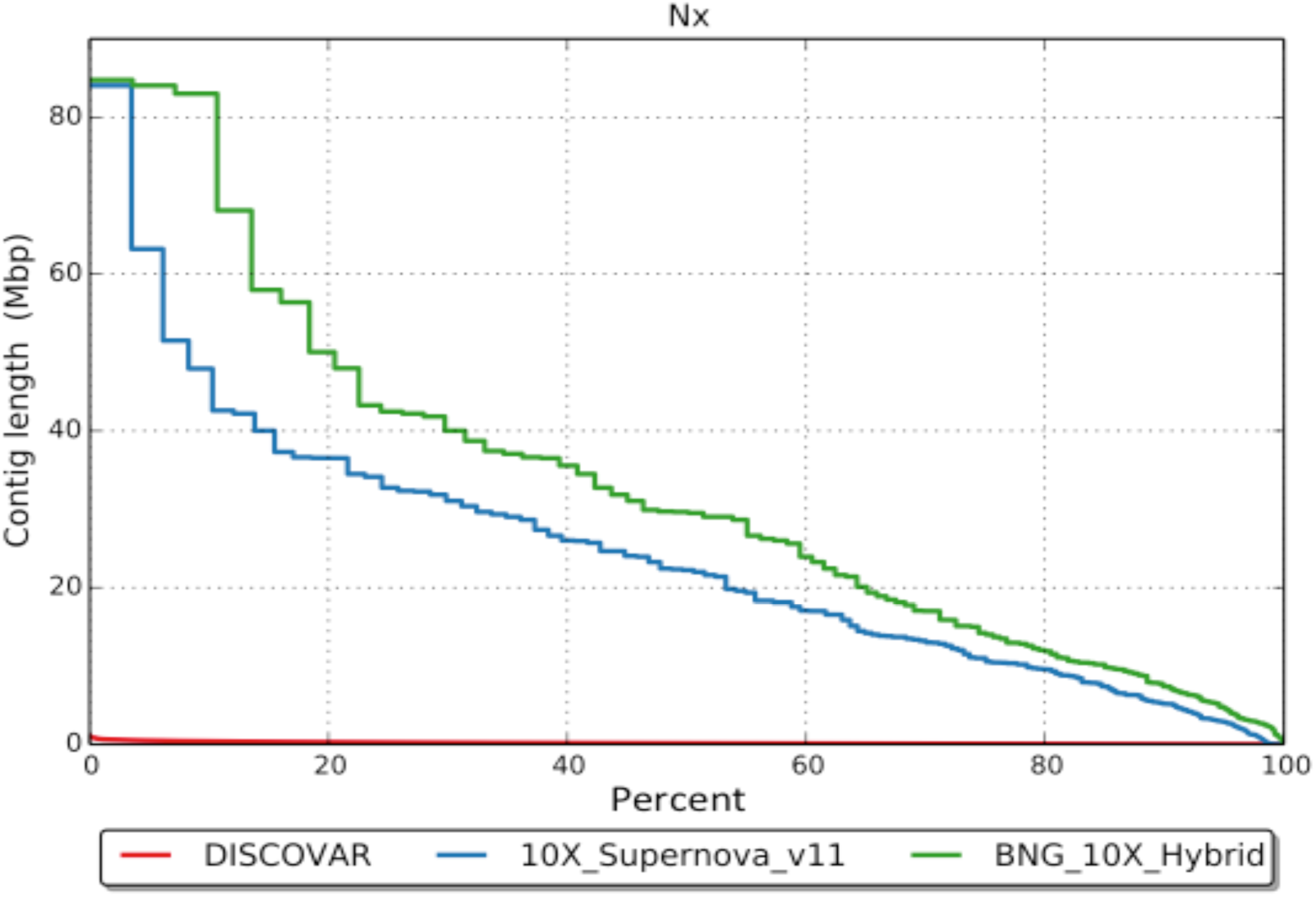
A QUAST (Gurevich et al., 2013) Nx plot of contig lengths as a percent of total scaffolds. The red line represents data assembled by DISCOVAR *de novo* (https://www.broadinstitute.org/software/discovar/blog/; Weisenfeld et al., 2014). Blue shows the improved scaffold lengths using the 10X Genomics Chromium chemistry and Supernova assembler v1.1. The green line is the additional improvement of the Supernova scaffolds when combined with Bionano Genomics optical maps.

**Fig 2.**
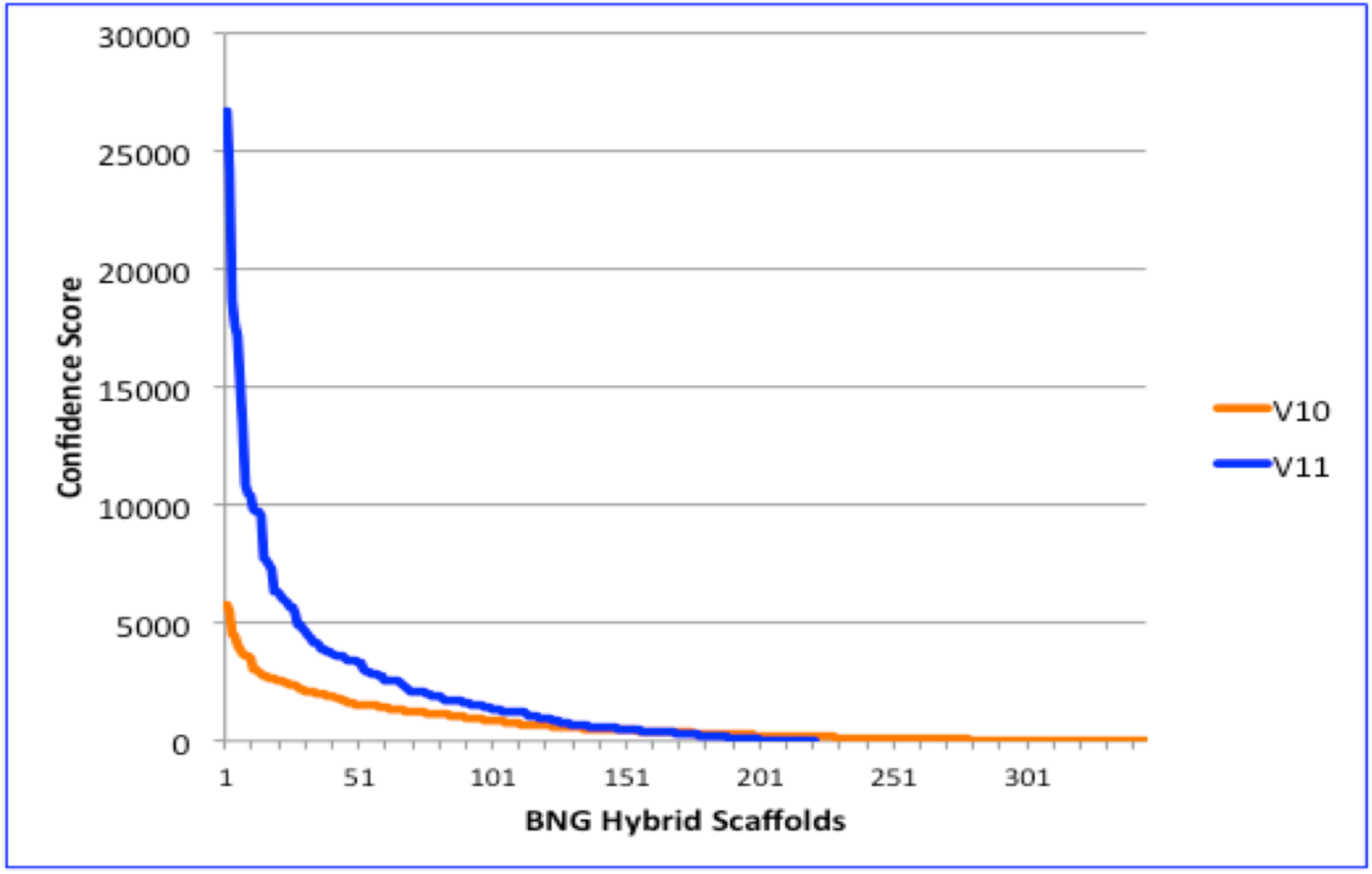
Improved gap estimation in Supernova v1.1 significantly improved the Bionano confidence scores while reducing the total number of hybrid scaffolds.

**Figure 3.**
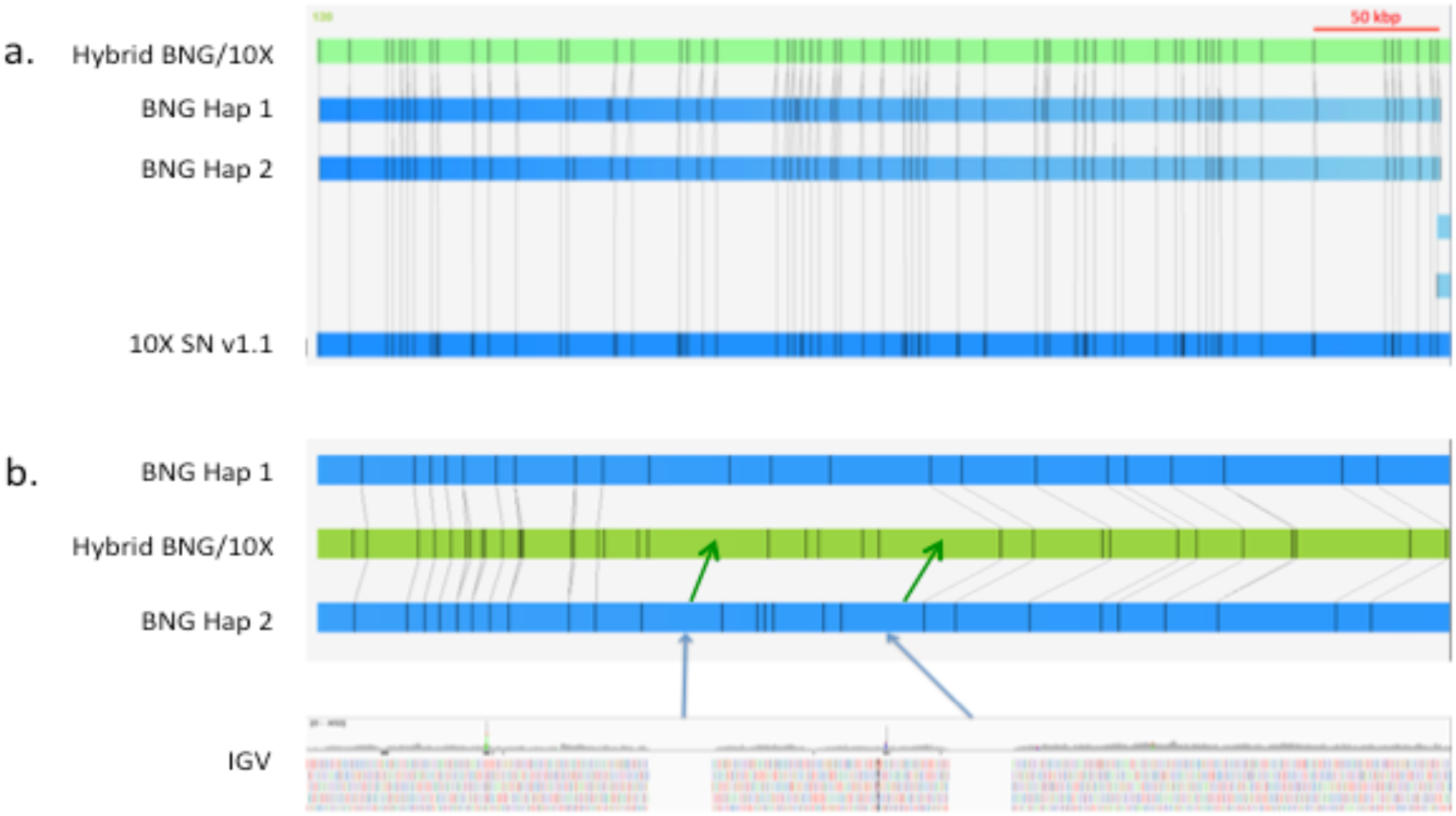
a. Example of a long block of high confidence alignment between Bionano optical maps and the Supernova v1.1 sequence. b. Example of where Supernova v1.1 (green arrows) estimated gap sizes that appeared larger than observed optically (blue arrows).

**Figure 4.**
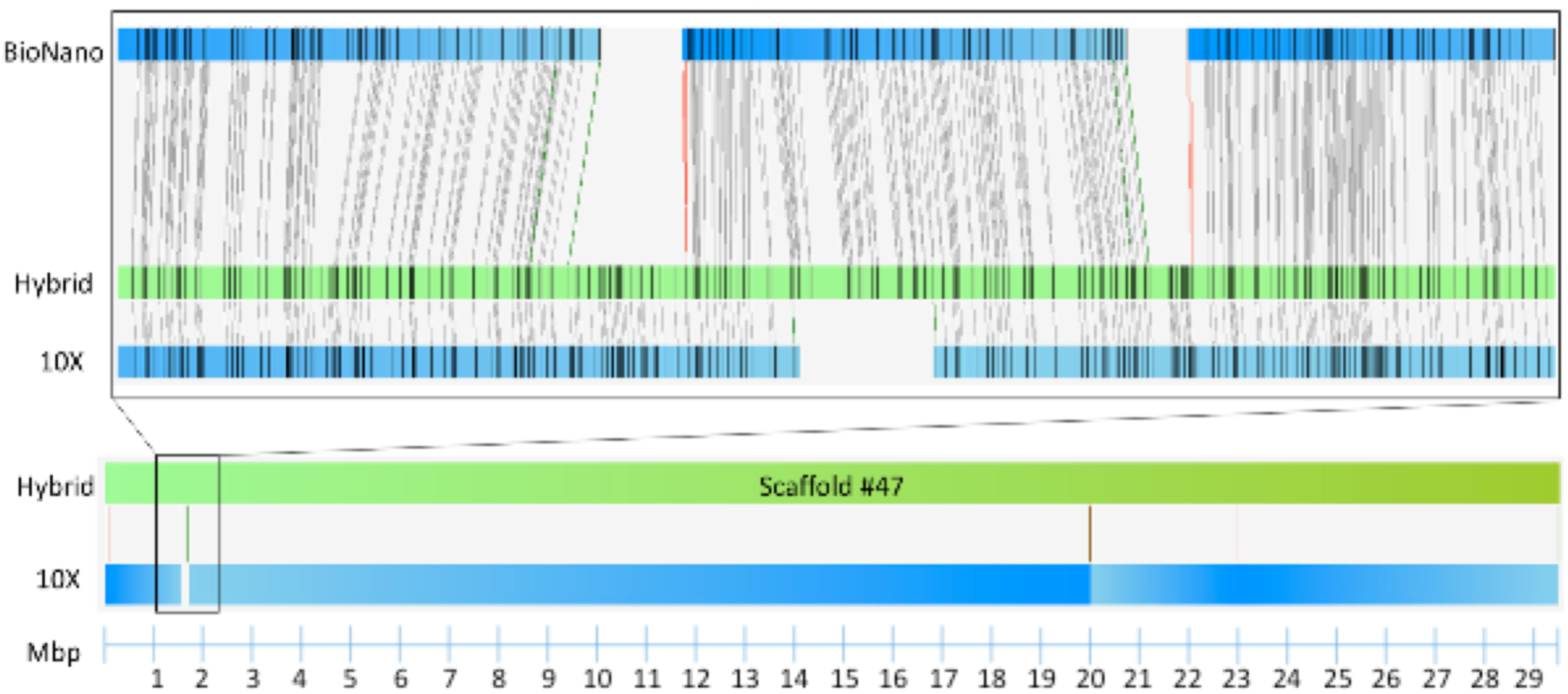
Bionano hybrid scaffold #47 with magnification, above, of the region at approximately 1.7 Mbp. This region is a position of evolutionary breakage or inversion in mammalian genomes. The optical maps confirmed that the DNA scaffolds were correctly joined.

In order to identify possible false Bionano joins between Supernova scaffolds, we performed TBLASTN, using a locally installed version of BLAST, to compare the 170 Bionano/10XG hybrid scaffolds against a database of human protein-coding genes. We set the match criteria at low stringency (evalue 1e-05) so that we would maximize the likelihood of identifying homologs but with the understanding that there would be many false matches. We inspected matches by finding two or more contiguous genes that mapped to the same human chromosome and with similar spacing and order. In a few instances of gene families with highly similar paralogs the exactly matching ortholog was not always identified by TBLASTN and in these cases we assigned the specific gene that best fit the conserved synteny relationship.

As an example, Table 2 shows four non-overlapping scaffolds that share conserved synteny with the majority of human chromosome 6. Although we noted inversions in gene order within scaffolds when they were arranged with respect to the chromosome we saw excellent coverage with no large gaps of expected orthologs. When the same analysis was extended to the remainder of the genome (data not shown) we found similar coverage with relatively few gaps indicating that most of the protein-coding genes have been captured in the 170 hybrid scaffolds. The only regions conspicuously absent were the short arms of the acrocentric human chromosome which are largely ribosomal RNA repeats and would not have been identified by TBLASTN analysis. The seal scaffolds orthologous to human chromosome 6 from Fig. 4 are shown in Fig. 5 as a nucmer plot which compares DNA sequences between seal and human. The plot confirms that the sequence contiguity within the seal scaffolds agrees with the conserved synteny analysis.

**Table 2.** Four seal scaffolds with genes orthologous to human chr 6. The orthologs were ordered according to their conserved syntenic positions on chr 6 and extended the length of the chromosome. The order of the genes highlighted in yellow in scaffold 47 was inverted relative to the human gene order. The inversion was confirmed by Bionano optical maps (Fig. 4).

| Scaffold | Position | RefGene | HuChr | Start | End |
| --- | --- | --- | --- | --- | --- |
| 47 | 2,841,008 | EXOC2 | 6 | 485,133 | 693,111 |
| 47 | 2,953,434 | HUS1B | 6 | 655,939 | 656,963 |
| 47 | 5,432,511 | MYLK4 | 6 | 2,663,629 | 2,750,966 |
| 47 | 5,637,247 | SERPINB9 | 6 | 2,887,266 | 2,903,280 |
| 47 | 6,850,365 | ECI2 | 6 | 4,115,689 | 4,135,597 |
| 47 | 7,644,508 | RPP40 | 6 | 4,994,732 | 5,004,063 |
| 47 | 10,025,315 | RIOK1 | 6 | 7,389,496 | 7,418,037 |
| 47 | 10,624,264 | BLOC1S5 | 6 | 8,013,567 | 8,064,414 |
| 47 | 11,007,567 | SLC35B3 | 6 | 8,413,068 | 8,435,483 |
| 47 | 13,360,854 | PAK1IP1 | 6 | 10,694,695 | 10,709,782 |
| 47 | 13,481,103 | GCM2 | 6 | 10,873,223 | 10,881,941 |
| 47 | 14,288,117 | ADTRP | 6 | 11,712,054 | 11,807,046 |
| 47 | 14,798,305 | EDN1 | 6 | 12,290,363 | 12,297,194 |
| 47 | 16,055,443 | RANBP9 | 6 | 13,621,498 | 13,711,564 |
| 47 | 16,211,441 | MCUR1 | 6 | 13,786,549 | 13,814,994 |
| 47 | 16,500,550 | CD83 | 6 | 14,117,256 | 14,136,918 |
| 47 | 17,589,062 | JARID2 | 6 | 15,246,296 | 15,522,040 |
| 47 | 18,282,073 | MYLIP | 6 | 16,129,125 | 16,148,248 |
| 47 | 19,523,903 | FAM8A1 | 6 | 17,600,355 | 17,611,719 |
| 47 | 20,011,483 | KDM1B | 6 | 18,155,329 | 18,223,853 |
| 47 | 22,093,462 | E2F3 | 6 | 20,401,906 | 20,493,715 |
| 47 | 25,658,069 | GPLD1 | 6 | 24,424,565 | 24,495,205 |
| 47 | 25,881,304 | C6orf62 | 6 | 24,704,861 | 24,720,836 |
| 47 | 26,768,511 | SLC17A4 | 6 | 25,754,699 | 25,781,191 |
| 47 | 27,610,706 | PRSS16 | 6 | 27,247,701 | 27,256,624 |
| 47 | 28,725,579 | GPX6 | 6 | 28,503,296 | 28,528,215 |
| 47 | 29,480,987 | GABBR1 | 6 | 29,602,228 | 29,633,135 |
| 47 | 2,917,479 | MOG | 6 | 29,656,981 | 29,672,372 |
| 47 | 2,252,098 | HLA-F | 6 | 29,723,340 | 29,740,355 |
| 47 | 2,251,182 | HLA-A | 6 | 29,942,470 | 29,945,884 |
| 47 | 2,108,886 | C6orf136 | 6 | 30,647,039 | 30,653,210 |
| 47 | 1,936,783 | DDR1 | 6 | 30,870,550 | 30,889,733 |
| 47 | 1,453,436 | DDX39B | 6 | 31,606,569 | 31,622,924 |
| 47 | 1,284,137 | CLIC1 | 6 | 31,730,581 | 31,737,318 |
| 47 | 1,026,563 | TNXB | 6 | 31,998,796 | 32,043,729 |
| 47 | 1,050,161 | CYP21A2 | 6 | 32,014,113 | 32,017,485 |
| 47 | 945,897 | PBX2 | 6 | 32,133,204 | 32,138,654 |
| 47 | 932,650 | NOTCH4 | 6 | 32,194,843 | 32,224,067 |
| 152 | 38,345 | HLA-DRB5 | 6 | 32,517,343 | 32,530,287 |
| 152 | 60,960 | HLA-DRB1 | 6 | 32,552,990 | 32,589,848 |
| 152 | 62,928 | HLA-DQB1 | 6 | 32,659,467 | 32,668,383 |
| 152 | 62,934 | HLA-DPB1 | 6 | 33,075,926 | 33,087,201 |
| 152 | 428,940 | COL11A2 | 6 | 33,162,681 | 33,192,499 |
| 152 | 447,468 | SLC39A7 | 6 | 33,200,445 | 33,204,439 |
| 152 | 506,118 | B3GALT4 | 6 | 33,277,132 | 33,284,832 |
| 152 | 610,185 | CUTA | 6 | 33,416,442 | 33,418,317 |
| 152 | 737,381 | BAK1 | 6 | 33,572,547 | 33,580,293 |
| 152 | 793,212 | ITPR3 | 6 | 33,620,365 | 33,696,574 |
| 152 | 848,284 | UQCC2 | 6 | 33,694,293 | 33,711,727 |
| 152 | 1,166,129 | GRM4 | 6 | 34,018,645 | 34,155,622 |
| 152 | 1,502,892 | RPS10 | 6 | 34,417,454 | 34,426,125 |
| 152 | 1,584,108 | PACSIN1 | 6 | 34,466,061 | 34,535,231 |
| 152 | 1,859,803 | UHRF1BP1 | 6 | 34,792,015 | 34,883,138 |
| 152 | 2,066,563 | TCP11 | 6 | 35,118,071 | 35,148,610 |
| 152 | 2,343,660 | PPARD | 6 | 35,342,558 | 35,428,191 |
| 152 | 2,369,881 | FANCE | 6 | 35,452,361 | 35,467,103 |
| 152 | 2,417,269 | TULP1 | 6 | 35,497,874 | 35,512,938 |
| 152 | 2,656,648 | ARMC12 | 6 | 35,737,032 | 35,749,079 |
| 152 | 2,744,242 | SRPK1 | 6 | 35,832,966 | 35,921,342 |
| 152 | 2,841,367 | SLC26A8 | 6 | 35,943,514 | 36,024,868 |
| 152 | 2,994,721 | MAPK14 | 6 | 36,027,677 | 36,111,236 |
| 152 | 3,088,997 | BRPF3 | 6 | 36,196,744 | 36,232,790 |
| 152 | 3,204,414 | C6orf222 | 6 | 36,315,757 | 36,336,885 |
| 152 | 3,348,241 | KCTD20 | 6 | 36,442,767 | 36,491,143 |
| 152 | 3,368,929 | STK38 | 6 | 36,493,892 | 36,547,470 |
| 152 | 3,577,284 | CPNE5 | 6 | 36,740,775 | 36,840,002 |
| 152 | 3,717,442 | C6orf89 | 6 | 36,871,870 | 36,928,964 |
| 152 | 3,827,924 | FGD2 | 6 | 37,005,646 | 37,029,070 |
| 152 | 4,126,093 | RNF8 | 6 | 37,353,972 | 37,394,738 |
| 152 | 4,231,592 | CMTR1 | 6 | 37,433,131 | 37,482,844 |
| 152 | 4,821,753 | ZFAND3 | 6 | 37,819,499 | 38,154,624 |
| 152 | 5,284,776 | BTBD9 | 6 | 38,168,451 | 38,640,148 |
| 152 | 5,363,014 | GLO1 | 6 | 38,675,925 | 38,703,141 |
| 152 | 5,418,483 | DNAH8 | 6 | 38,715,341 | 39,030,529 |
| 152 | 5,810,207 | GLP1R | 6 | 39,048,798 | 39,087,743 |
| 152 | 6,024,324 | KCNK16 | 6 | 39,314,698 | 39,322,968 |
| 152 | 6,047,898 | KIF6 | 6 | 39,329,990 | 39,725,405 |
| 26 | 406,011 | DAAM2 | 6 | 39,792,298 | 39,904,877 |
| 26 | 2,001,969 | FOXP4 | 6 | 41,546,426 | 41,602,384 |
| 26 | 2,206,234 | USP49 | 6 | 41,789,896 | 41,895,361 |
| 26 | 2,408,430 | TAF8 | 6 | 42,050,513 | 42,087,461 |
| 26 | 2,445,724 | C6orf132 | 6 | 42,101,118 | 42,142,619 |
| 26 | 2,508,153 | GUCA1B | 6 | 42,184,401 | 42,194,916 |
| 26 | 2,892,519 | UBR2 | 6 | 42,564,062 | 42,693,504 |
| 26 | 3,876,392 | VEGFA | 6 | 43,770,184 | 43,786,487 |
| 26 | 4,372,743 | AARS2 | 6 | 44,299,654 | 44,313,326 |
| 26 | 6,042,112 | ENPP4 | 6 | 46,129,993 | 46,146,699 |
| 26 | 7,254,492 | CD2AP | 6 | 47,477,789 | 47,627,263 |
| 26 | 9,063,622 | MUT | 6 | 49,430,360 | 49,463,191 |
| 26 | 10,266,365 | TFAP2D | 6 | 50,713,828 | 50,772,988 |
| 26 | 10,345,803 | TFAP2B | 6 | 50,818,723 | 50,847,613 |
| 26 | 11,729,609 | MCM3 | 6 | 52,264,009 | 52,284,881 |
| 26 | 13,399,950 | MLIP | 6 | 53,929,982 | 54,266,280 |
| 26 | 14,055,747 | FAM83B | 6 | 54,846,771 | 54,942,022 |
| 26 | 14,562,753 | HMGCLL1 | 6 | 55,434,369 | 55,579,214 |
| 26 | 15,552,546 | DST | 6 | 56,457,987 | 56,954,628 |
| 26 | 16,056,305 | BEND6 | 6 | 56,955,126 | 57,027,342 |
| 26 | 16,216,175 | BAG2 | 6 | 57,172,326 | 57,189,833 |
| 26 | 16,364,502 | PRIM2 | 6 | 57,314,805 | 57,646,849 |
| 26 | 18,733,835 | LGSN | 6 | 63,275,951 | 63,319,977 |
| 26 | 19,020,962 | PHF3 | 6 | 63,635,820 | 63,779,336 |
| 26 | 23,830,984 | AGRB3 | 6 | 68,635,282 | 69,389,511 |
| 26 | 25,018,666 | FAM135A | 6 | 70,412,941 | 70,561,174 |
| 26 | 25,658,762 | OGFRL1 | 6 | 71,288,803 | 71,308,950 |
| 3 | 84,112,823 | RIMS1 | 6 | 71,886,703 | 72,403,143 |
| 3 | 83,186,090 | MTO1 | 6 | 73,461,578 | 73,509,236 |
| 3 | 81,949,665 | COL12A1 | 6 | 75,084,326 | 75,206,051 |
| 3 | 81,198,935 | IMPG1 | 6 | 75,921,115 | 76,072,678 |
| 3 | 78,875,241 | IRAK1BP1 | 6 | 78,867,472 | 78,946,440 |
| 3 | 78,731,298 | PHIP | 6 | 78,935,867 | 79,078,236 |
| 3 | 77,882,873 | ELOVL4 | 6 | 79,914,812 | 79,947,580 |
| 3 | 76,413,900 | FAM46A | 6 | 81,491,439 | 81,752,774 |
| 3 | 75,114,258 | DOPEY1 | 6 | 83,067,666 | 83,171,350 |
| 3 | 75,043,295 | PGM3 | 6 | 83,161,150 | 83,193,936 |
| 3 | 74,977,755 | ME1 | 6 | 83,210,389 | 83,431,071 |
| 3 | 71,714,453 | C6orf163 | 6 | 87,344,849 | 87,365,463 |
| 3 | 71,605,411 | SLC35A1 | 6 | 87,470,623 | 87,512,336 |
| 3 | 71,564,591 | RARS2 | 6 | 87,514,378 | 87,590,003 |
| 3 | 70,176,222 | PM20D2 | 6 | 89,146,050 | 89,165,565 |
| 3 | 69,625,479 | MDN1 | 6 | 89,642,499 | 89,819,723 |
| 3 | 66,247,239 | EPHA7 | 6 | 93,240,020 | 93,419,547 |
| 3 | 64,349,305 | MANEA | 6 | 95,577,543 | 95,609,457 |
| 3 | 63,458,934 | UFL1 | 6 | 96,521,595 | 96,555,276 |
| 3 | 63,160,649 | NDUFAF4 | 6 | 96,889,313 | 96,897,881 |
| 3 | 60,825,374 | FAXC | 6 | 99,271,169 | 99,350,062 |
| 3 | 59,606,368 | ASCC3 | 6 | 100,508,194 | 100,881,372 |
| 3 | 58,505,231 | GRIK2 | 6 | 101,398,788 | 102,070,083 |
| 3 | 55,916,988 | HACE1 | 6 | 104,728,093 | 104,859,919 |
| 3 | 55,602,860 | POPDC3 | 6 | 105,158,280 | 105,179,995 |
| 3 | 55,407,039 | PREP | 6 | 105,277,565 | 105,403,084 |
| 3 | 54,230,441 | QRSL1 | 6 | 106,629,578 | 106,668,417 |
| 3 | 53,609,305 | PDSS2 | 6 | 107,152,557 | 107,459,683 |
| 3 | 53,229,894 | SEC63 | 6 | 107,867,756 | 107,958,278 |
| 3 | 53,324,808 | SCML4 | 6 | 107,704,104 | 107,824,317 |
| 3 | 52,662,962 | LACE1 | 6 | 108,294,874 | 108,525,784 |
| 3 | 52,242,062 | ARMC2 | 6 | 108,848,416 | 108,974,472 |
| 3 | 51,792,347 | PPIL6 | 6 | 109,390,215 | 109,441,171 |
| 3 | 51,555,610 | FIG4 | 6 | 109,691,312 | 109,825,428 |
| 3 | 50,548,418 | AMD1 | 6 | 110,874,770 | 110,895,713 |
| 3 | 49,349,830 | LAMA4 | 6 | 112,108,760 | 112,254,939 |
| 3 | 46,185,569 | FRK | 6 | 115,931,149 | 116,060,758 |
| 3 | 45,959,041 | NT5DC1 | 6 | 116,100,849 | 116,249,497 |
| 3 | 45,663,548 | FAM26F | 6 | 116,461,370 | 116,463,779 |
| 3 | 44,966,229 | ROS1 | 6 | 117,288,300 | 117,425,855 |
| 3 | 43,425,386 | MAN1A1 | 6 | 119,177,209 | 119,349,761 |
| 3 | 40,748,527 | HSF2 | 6 | 122,399,546 | 122,433,119 |
| 3 | 40,715,709 | SERINC1 | 6 | 122,443,354 | 122,471,822 |
| 3 | 40,445,056 | SMPDL3A | 6 | 122,789,049 | 122,809,720 |
| 3 | 37,577,092 | TRMT11 | 6 | 125,986,430 | 126,039,276 |
| 3 | 37,313,537 | CENPW | 6 | 126,340,174 | 126,348,875 |
| 3 | 36,442,589 | ECHDC1 | 6 | 127,288,710 | 127,343,609 |
| 3 | 34,410,632 | LAMA2 | 6 | 128,883,141 | 129,516,569 |
| 3 | 33,701,892 | SAMD3 | 6 | 130,144,315 | 130,365,425 |
| 3 | 32,529,868 | MED23 | 6 | 131,573,966 | 131,628,229 |
| 3 | 31,526,312 | VNN3 | 6 | 132,722,787 | 132,734,765 |
| 3 | 31,491,481 | SLC18B1 | 6 | 132,769,370 | 132,798,553 |
| 3 | 30,216,128 | SGK1 | 6 | 134,169,246 | 134,318,112 |
| 3 | 29,668,501 | ALDH8A1 | 6 | 134,917,390 | 134,950,122 |
| 3 | 29,578,641 | HBS1L | 6 | 134,960,378 | 135,054,898 |
| 3 | 29,426,667 | MYB | 6 | 135,181,315 | 135,219,173 |
| 3 | 28,465,143 | BCLAF1 | 6 | 136,256,863 | 136,289,851 |
| 3 | 27,841,899 | IL20RA | 6 | 136,999,971 | 137,045,180 |
| 3 | 27,111,169 | TNFAIP3 | 6 | 137,867,188 | 137,883,312 |
| 3 | 26,723,014 | ARFGEF3 | 6 | 138,161,921 | 138,344,663 |
| 3 | 25,844,008 | CITED2 | 6 | 139,371,807 | 139,374,620 |
| 3 | 23,556,896 | NMBR | 6 | 142,058,330 | 142,088,799 |
| 3 | 22,460,142 | AIG1 | 6 | 143,060,496 | 143,340,304 |
| 3 | 22,307,461 | PEX3 | 6 | 143,450,807 | 143,490,010 |
| 3 | 21,913,036 | PLAGL1 | 6 | 143,940,300 | 144,064,599 |
| 3 | 21,139,050 | UTRN | 6 | 144,285,701 | 144,853,034 |
| 3 | 20,188,841 | SHPRH | 6 | 145,864,245 | 145,964,423 |
| 3 | 19,496,651 | ADGB | 6 | 146,598,965 | 146,815,462 |
| 3 | 18,802,872 | SAMD5 | 6 | 147,508,927 | 147,737,547 |
| 3 | 17,136,380 | TAB2 | 6 | 149,218,641 | 149,411,613 |
| 3 | 16,889,963 | LATS1 | 6 | 149,658,153 | 149,718,256 |
| 3 | 15,547,830 | ARMT1 | 6 | 151,452,258 | 151,470,101 |
| 3 | 14,653,210 | SYNE1 | 6 | 152,121,684 | 152,637,801 |
| 3 | 14,217,199 | MTRF1L | 6 | 152,987,362 | 153,002,685 |
| 3 | 12,161,992 | NOX3 | 6 | 155,395,370 | 155,455,903 |
| 3 | 9,662,930 | SERAC1 | 6 | 158,109,515 | 158,168,270 |
| 3 | 8,290,345 | TCP1 | 6 | 159,778,498 | 159,789,703 |
| 3 | 8,061,191 | IGF2R | 6 | 159,969,099 | 160,113,507 |
| 3 | 7,775,898 | PLG | 6 | 160,702,238 | 160,753,315 |
| 3 | 5,807,350 | PACRG | 6 | 162,727,132 | 163,315,492 |
| 3 | 3,910,419 | C6orf118 | 6 | 165,279,664 | 165,309,607 |
| 3 | 2,290,679 | UNC93A | 6 | 167,271,169 | 167,316,019 |
| 3 | 960,211 | THBS2 | 6 | 169,215,780 | 169,254,044 |
| 3 | 39,405 | PSMB1 | 6 | 170,535,117 | 170,553,341 |
| 3 | 5,415 | TBP | 6 | 170,554,302 | 170,572,870 |

**Fig 5.**
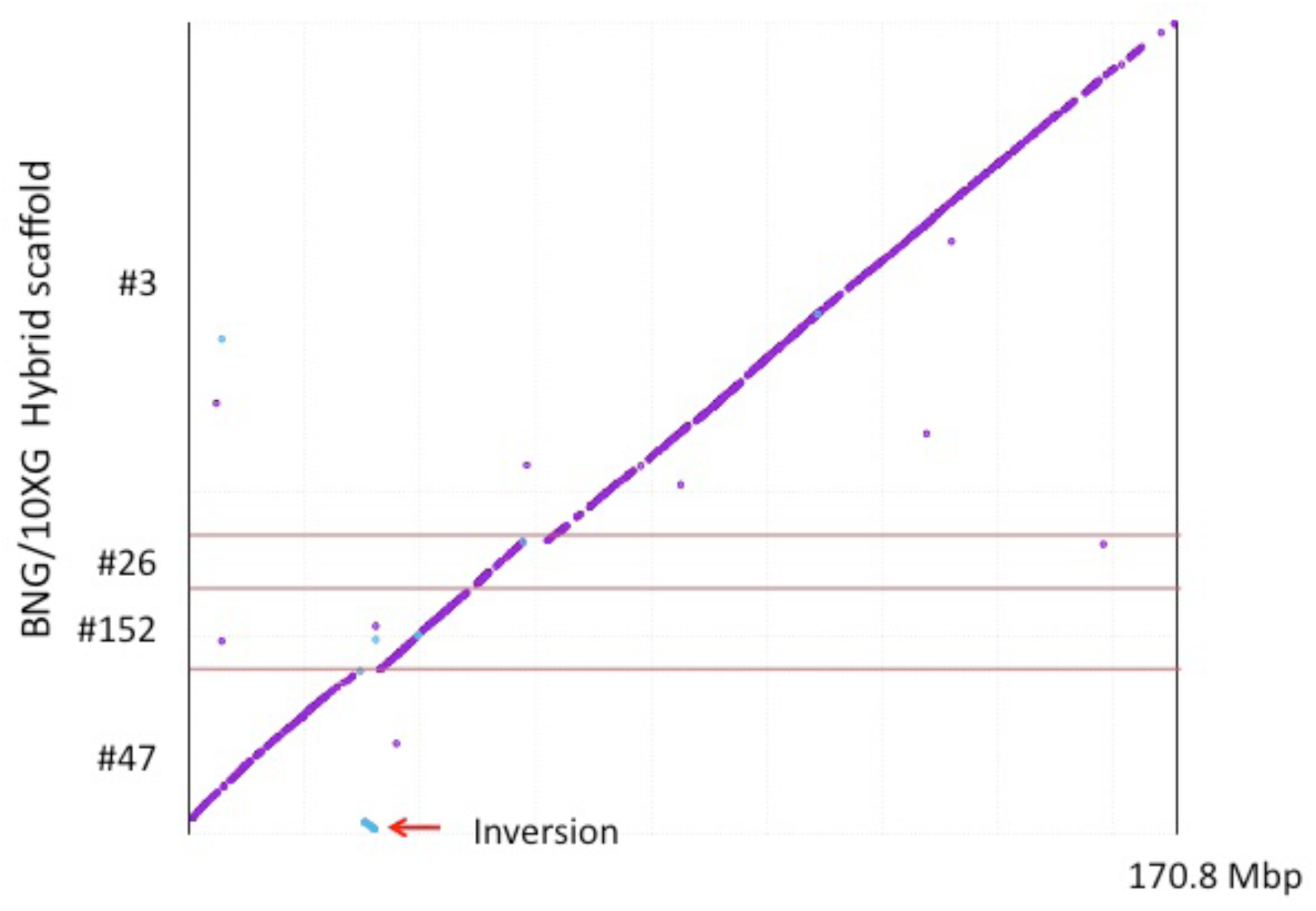
DNA alignment of the four scaffolds identified by TBLASTN conserved synteny analysis between the monk seal and human chromosome 6 (170.8 Mbp long). Inverted genes in Table 2 are indicated by the arrow.

We also performed BUSCO analysis of the conserved vertebrate gene set both for the 170 hybrid scaffolds as well as the unscaffolded sequences that did not align to Bionano maps. Of the 170 hybrid scaffolds BUSCO identified 2,696 complete single-copy genes (89.2% of the 3023 expected conserved vertebrate genes) and 16 duplicates (0.5%). These BUSCO assessments are nearly identical to those calculated for humans (Simão et al., 2015). An additional 171 genes were scored as fragmented (5.7%) and 140 were missing (4.6%). We manually searched our TBLASTN predicted protein list and accounted for all but 12 of the “missing.” Among the sequences not aligning to optical maps, 8 “complete” and 14 “fragmented” genes were found (fewer than 1% of the total screened).

## DISCUSSION

Next generation sequencing has revolutionized the field of comparative genomics, especially at the level of individual gene sequences. But, creating long assemblies has been less successful because of the complexity of genomes which are frequently interrupted by repeats longer than typical NGS reads. This limitation has led to various innovations such as long-single molecule sequencing (e.g., PacBio, Oxford Nanopore) but throughput, accuracy and costs have limited their widespread adoption. We showed that the 10XG Chromium method, assembled with Supernova v1.1 assembly software, produced long scaffolds 160-fold longer than paired end sequencing assembled with DISCOVAR *de novo*. When combined with Bionano optical maps the scaffold N50 increased to greater than 29 Mbp, a 200-fold improvement. The technologies described here are a large step toward improving and simplifying *de novo* genome assembly in that they use orthogonal methods that complement each other. The strengths of the 10XG Chromium Linked-Reads method are that it uses ∼ 106 unique molecular indices to randomly tag long molecules captured in emulsified droplets and then sequences these by standard short-read technology. Because the total amount of DNA is so small (1.2 ng for a mammalian genome), the probability that more than one molecule from the same region of the genome is encapsulated and identically labeled is extremely small, and long molecules will produce linked reads that are likely to span repeats such as L1 and other retrotransposons.

Dong et al. (2013) used optical mapping to assist with the assembly of the goat genome and found that it improved super-scaffold lengths by 5X over those obtained with fosmid end sequencing. Mostovoy et al. (2016) used 10XG chemistry and Bionano Genomics maps to create a phased genome assembly of the widely sequenced human sample NA12878 and reported 170 hybrid scaffolds with an N50 of 33.5 Mbp. We have extended their approach by using newer versions of the 10XG chemistry with an increased number of UMIs and Supernova software for *de novo* assembly. Although optical maps do not produce sequence *per se* they do provide a valuable “truth set” of chromosomal structure to confirm or refute sequence assemblies. The Bionano optical maps were useful both for merging 10X scaffolds but also as a quality assessment measure of the overall sequence contiguity in any given scaffold. Although the N50 statistic can be misleading based on the choice of the lower contig cutoff size, the assembly that we produced by combining the 10XG and Bionano optical maps resulted in larger scaffolds than most mammalian genomes reported to date. The hybrid Bionano/10XG scaffolds also had conserved syntenic blocks that were consistent with the human genome and other mammals. The QUAST statistics (Table 1) indicated that 98.3% of the total estimated genome length (i.e., the predicted haploid genome equivalent; 2360/2400 Mbp) was accounted for by the 170 Bionano hybrid scaffolds. This is in agreement with the conserved synteny data which showed that most human chromosome orthologous regions were very well covered. As noted above, BUSCO identified 99% of the conserved gene set within the 170 scaffolds, again indicating that the hybrid scaffolds were of high quality.

The materials cost for this study was less than $15,000. We anticipate that these costs will become lower in the near term as optical mapping improves, short-read sequencing migrates to higher throughput instruments, or new long read technologies (e.g., nanopore sequencing) mature. Software improvements and more standardized pipelines for genome assembly are also expected to make genome assembly less burdensome.

## CONCLUSION

As a field, comparative genomics is entering a new phase where we expect that the vast majority of living (or recently extinct) species will be sequenced. We know from human studies that many of the regulatory signals that control complex traits and development are in intergenic regions (e.g., Bhatia and Kleinjan, 2014; GWAS Catalog: http://www.ebi.ac.uk/gwas/). Likewise, both the sequences and the genomic architecture of regulatory elements within chromosomes are likely to explain the majority of the phenotypic differences between species. If that is true then a fuller understanding of comparative genomics will require knowing not only selected gene sequences but the context of the genome in which those genes are found.

Combining the two orthogonal methods of Chromium Linked-Reads with Bionano optical maps is likely to make the assembly of high quality genomes routine and significantly improve our understanding of comparative genome biology while reducing costs. Because the Chromium method labels single molecules only a few ng of DNA are needed. This advantage may be particularly useful for non-lethal sampling of organisms both in captivity and in the field. As more species become endangered the need to preserve their genomes for study and conservation will increase. The methods shown here represent a cost-effective and robust strategy to meet that goal.

## Acknowledgements

We thank Drs. Charles Littnan and Michelle Barbieri for obtaining samples, Dr. Susan Brown for performing the optical mapping, Dr. Melissa Olson for lymphocyte separation and whole blood cryopreservation, Laura Kasch and Jill Barton for preparing agarose blocks for optical mapping and DNA isolation from blood, Drs. Jonathan Pevsner and Michael Schatz for comments on the manuscript.

## Supplementary materials

### Analysis Commands

BLAST:

~~~
tblastn -num_descriptions 10000 -num_alignments 10000 -evalue 1e-05 -outfmt “7 qacc sacc evalue qstart qend sstart send length score stitle”
~~~

BIONANO Hybrid Assembly:

~~~
perl /home/bionano/scripts/HybridScaffold/hybridScaffold.pl -n
~~~

~~~
_fastasassembly_pseudohap2.1.fasta -b EXP_REFINEFINAL1.cmap -c
~~~

~~~
/home/bionano/scripts/HybridScaffold/hybridScaffold_config_aggressive.xml -o
~~~

~~~
./hybridOutput1_noCut -r /home/bionano/tools/RefAligner -B 2 -N 1 -f
~~~

BUSCOv2:

~~~
BUSCO.py –i reference.fasta –o output_name –l vertebrata -m genome -c 4 -sp human
~~~

DISCOVARdenovo:

~~~
DiscovarDeNovo READS=“ *.fastq.gz” NUM_THREADS=48 MAX_MEM_GB=900 OUT_DIR=my_discovar_assembly
~~~

NUCMER:

~~~
nucmer --maxmatch -c 20 -l 50 reference.fasta query.fasta delta-filter -l 1000 input.delta > out.delta
~~~

~~~
mummerplot -postscript –R reference.fasta –Q query.fasta -f –p out.delta
~~~

QUAST:

~~~
quast reference.fasta –o output_directory -t4 -m0 –l my_label contig-thresholds 0,100,1000,10000,100000,1000000,10000000
~~~

SUPERNOVA:

~~~
supernova run –fastqs=demultiplexed_fastq_path --lanes=1,2
~~~

~~~
--indices=ATTCCGATA,ATTCCGATC,CCCTAACAA,CCTAACAAT,GAAGGCTGA,TGGATTGCA
~~~

~~~
--maxreads=975000000 --id=benny --description=monk_seal
~~~

